# Codon Bias Confers Stability to mRNAs via ILF2 in Humans

**DOI:** 10.1101/585992

**Authors:** Fabian Hia, Sheng Fan Yang, Yuichi Shichino, Masanori Yoshinaga, Yasuhiro Murakawa, Alexis Vandenbon, Akira Fukao, Toshinobu Fujiwara, Markus Landthaler, Tohru Natsume, Shungo Adachi, Shintaro Iwasaki, Osamu Takeuchi

## Abstract

Codon bias has been implicated as one of the major factors contributing to mRNA stability in yeast. However, the effects of codon-bias on mRNA stability remain unclear in humans. Here we show that human cells possess a mechanism to modulate RNA stability through a unique codon bias different from that of yeast. Bioinformatics analysis showed that codons could be clustered into two distinct groups – codons with G or C at the third base position (GC3) and codons with either A or T at the third base position (AT3); the former stabilizing while the latter destabilizing mRNA. Quantification of codon bias showed that increased GC3 content entails proportionately higher GC content. Through bioinformatics, ribosome profiling and *in vitro* analysis, we show that decoupling of the effects of codon bias reveals two modes of mRNA regulation, GC3- and GC-content dependent. Employing an immunoprecipitation-based strategy, we identified ILF2 as an RNA binding protein that differentially regulates global mRNA abundances based on codon bias. Our results demonstrate that codon bias is a two-pronged system that governs mRNA abundance.

## Introduction

Messenger RNA (mRNA) regulation represents an essential part of regulating a myriad of physiological processes in cells, being indicated in the maintenance of cellular homeostasis to immune responses [1–3]. In addition to transcription regulation, post-transcriptional regulation of mRNA stability is vital to the fine-tuning of mRNA abundance. To date, several mRNA-intrinsic properties, often in 5′ or 3′ untranslated regions (UTR), have been shown to affect mRNA stability [4,5]. Due to the recent advances in technology, the contribution of mRNA stability to gene expression has been suggested [6]. However, the regulation of mRNA stability, which is possibly governed by mRNA intrinsic features, has not been fully elucidated.

One of the most crucial mRNA-intrinsic features is codon bias. To scrutinize this bias in usage of redundant codons, several metrics to measure how efficiently codons are decoded by ribosomes (codon optimality) have been proposed. In a classical metric called the codon Adaptation Index (cAI), gene optimality is calculated by comparison between codon usage bias of a target gene and reference genes which are highly expressed [7,8]. Another index termed the tRNA Adaption Index (tAI) gauges how efficiently tRNA is utilized by the translating ribosome [9,10]. More recently, the normalized translation efficiency (nTE), which takes into consideration not only the availability of tRNA but also demand, was also proposed [11].

Recently, Presnyak and colleagues showed that mRNA half-lives are correlated with optimal codon content based on a metric, the Codon Stabilization Coefficient (CSC) which was calculated from the correlations between the codon frequencies in mRNAs and stabilities of mRNAs. Additionally, they showed that the substitutions of codons with their synonymous optimal and non-optimal counterparts resulted in significant increases and decreases in mRNA stability in yeast [12]. This effect was brought by an RNA binding protein (RBP) Dhh1p (mammalian ortholog DDX6), which senses ribosome elongation speed [12–14]. In yeast, these differences in ribosome elongation speed in turn are influenced by tRNA availability and demand [11,15,16]. Taken together, codons can be designated into optimal and non-optimal categories; the former hypothesized to be decoded efficiently and accurately [17,18] while the latter slow ribosome elongation resulting in decreased mRNA stability [12–14]. It is also important to make the distinction that common and rare codons do not necessarily imply optimal and non-optimal codons.

At present, codon optimality-mediated decay has been extensively studied and established particularly in *Saccharomyces cerevisiae* as well as other model organisms such as *Schizosaccharomyces pombe*, *Drosophila melanogaster*, *Danio rerio*, *Escherichia coli*, and *Neurospora crassa* [19–23]. Nevertheless to date, this system of codon optimality has been inadequately scrutinized in humans.

In this study, we show that codon bias-mediated decay exists in humans. Principal component analysis (PCA) showed that codons could be clustered into two distinct groups; codons with A or T at the third base position (AT3) and codons with either G or C at the third base position (GC3). This clustering was associated with mRNA half-lives enabling us to determine GC3 and AT3 codons as stabilizing and non-stabilizing codons respectively. In this regard, the increased usage of GC3 codons entails an inevitable increase GC-content. We then developed an algorithm to quantify the codon bias of GC3 codons. With ribosome profiling, we show that codon bias-derived occupancy scores agreed with ribosome occupancy. Additionally, bioinformatics analysis revealed that frameshifts abrogate this GC3-AT3 delineation. We then verified our results *in vitro* using optimized and de-optimized reporter constructs. Here we propose that GC3 codons and AT3 codons are optimized and de-optimized codons respectively. Importantly, frameshifted optimized transcripts retain a certain level of stability suggesting that overall the overall GC content of transcripts is an additional determinant of stability. Finally, employing a ribonucleoprotein immunoprecipitation strategy, we identified RNA binding proteins which were bound to transcripts with low or high GC3-content. We propose that interleukin enhancer-binding factor 2 (ILF2) mediates mRNA stability of transcripts via codon bias.

## Results

### Codons in *Homo sapiens* can be categorized into GC3 and AT3 codons

To examine whether a system of codon bias exists in human, we first compared codon frequencies in *Homo sapiens* and other model organisms. Hierarchical clustering analysis of codon frequency data obtained from Ensembl database [24] showed a difference between lower eukaryotes such as *Saccharomyces cerevisiae* and *Caenorhabditis elegans*, and higher eukaryotes such as *Homo sapiens* and *Mus musculus* (Fig 1A). To investigate codon bias in humans, we then calculated the codon frequencies of individual genes from *H. sapiens* and performed a principal component analysis (PCA) on the data. The first principal component (PC1) of the PCA which accounted for 22.85% of the total variance, divided codons into two clusters: codons with either G or C at the third base position (GC3) and codons with either A or T at the third base position (AT3) (Fig 1B). Interestingly, the division within the second principal component (PC2) appeared to be split along the number of G/C or A/T bases in codons. We repeated our analysis on the CDS sequences from *S. cerevisiae* and found no such clustering (**Fig EV1A**). However, we discovered that the factor loading scores of the codons along the first principal component of our analysis in yeast corresponded to the CSC metric [12], albeit differences in the order (**Fig EV1B**). The above-mentioned results therefore raised the possibility that GC3 and AT3 codons in humans may have a valid effect on mRNA stability.

**Figure 1.**
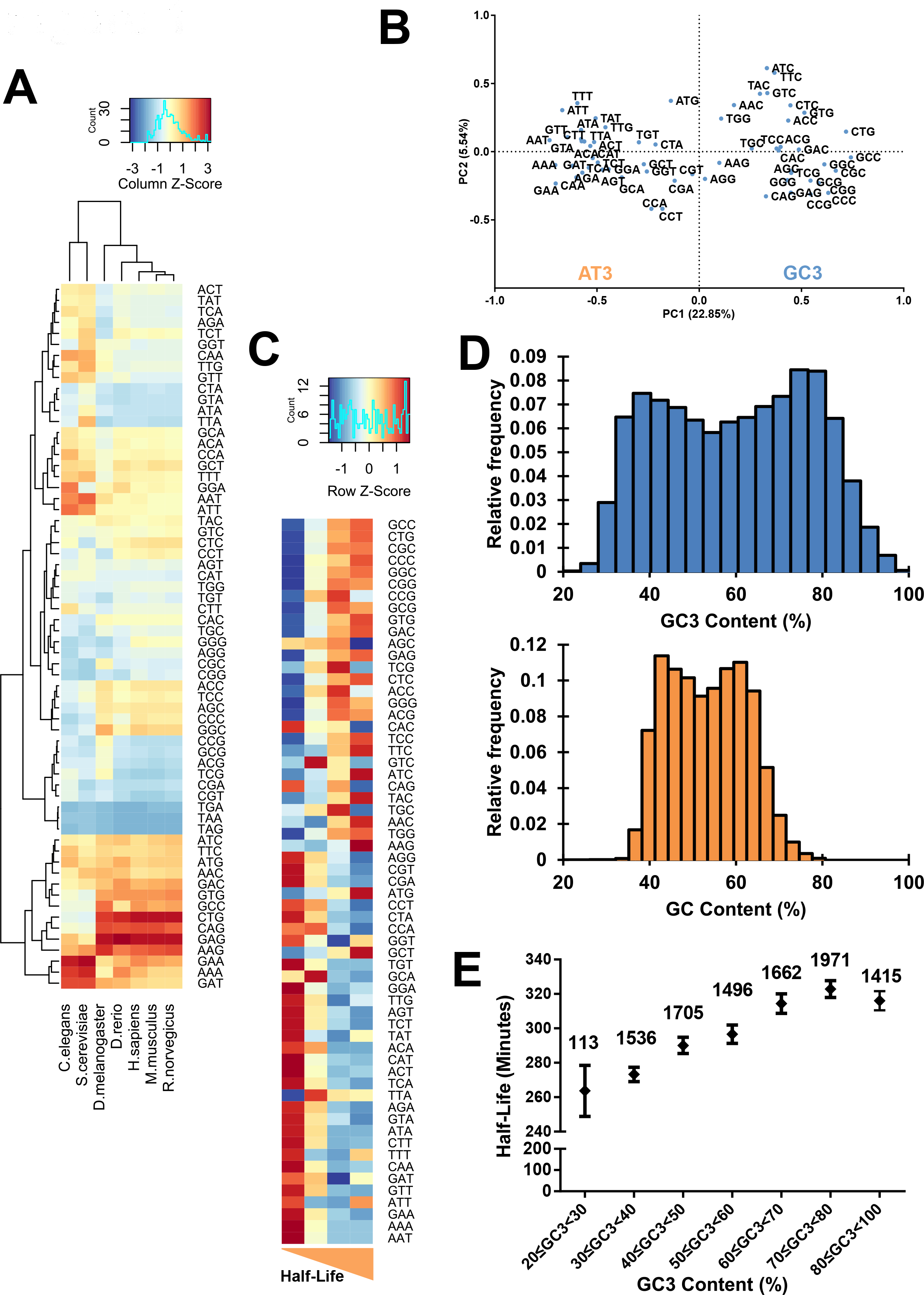
Bioinformatics analysis reveals that biased codons can be categorized into GC3 and AT3 codons respectively. **A.** Hierarchical clustering analysis of model organisms and their average CDS codon frequencies. **B.** Principal component analysis of the CDS codon frequencies of 9666 protein-coding genes. PC1 and PC2 indicate the first and second principal components. **C.** Heatmap of half-lives of mRNA and their CDS codon frequencies. The transcripts were ranked according to their half-lives and divided equally into quartiles. The respective codon frequencies of each group were then averaged. **D.** Histogram illustrating the distribution of genes and their respective GC3- and GC-content. **E.** Comparison of average transcript mRNA half-lives across their respective GC3-content ranges. Number of transcripts within each gene optimality range is indicated above their respective points. **Data information:** In (**E**), error bars represent the 95% confidence intervals.

We tested the link between mRNA stability and GC3-AT3 codons using published datasets of global mRNA decay rates in physiologically growing HEK293 cells (GSE69153) [25; Data ref: Murakawa et al, 2015]. Briefly, we divided the transcripts equally into quartiles based on their half-lives and averaged the codon frequencies within the quartiles. Strikingly, genes with short half-lives were associated with AT3 codons while genes with longer half-lives were associated with GC3 codons (Fig 1C), suggesting a connection between third base of codons and the stability of mRNAs.

Broadly, the codon bias in mRNA can predict the stability of the mRNA. By summing the GC3 frequencies and GC bases of CDS sequences, we could determine the GC3- and GC- content of a gene (**Dataset EV1**). We then visualized the GC3 and GC landscape by plotting the the corresponding values via a histogram (Figure 1D). GC3-content was represented as a bimodal distribution with a range of values from the minimum of 24.1% to the maximum of 100% while GC-content appeared similarly as a bimodal distribution with a range of values from a minimum of 27.6% to the maximum of 79.7%. A Pearson correlation analysis (R^2^ = 0.869) between gene GC-content and GC3-content (**Fig EV1C**) reflected an enrichment of GC-content with increased GC3-content. Indeed, higher GC3-content was generally associated with better stability (Fig 1E, **Fig EV1D**). Additionally, we noted that the codon bias *per se* was different between yeast and humans (Fig 1B **and Fig EV1A**) [12]. Taken together, our analysis allowed us to designate GC3 and AT3 codons as stabilizing and destabilizing codons respectively. Additionally, high GC3 content in transcripts inevitably results in high GC-content, which is a feature of stable mRNAs.

We then asked about the biological relevance associated with codon bias. Taking the 5% of lowest and highest ranked genes into account, we observed that genes with high GC3-content were enriched in developmental processes while genes with low GC3-content were enriched in cellular division processes (**Fig EV1E** and F), suggesting the importance of codon bias-mediated mRNA decay across dynamic cellular processes in humans.

### Ribosome profiling reveals that ribosome occupancy is correlated with codon bias

Given that GC3-AT3 codons were associated with high and low stability respectively, we wondered if these two groups were synonymous with optimal and non-optimal codons. It has been proposed that slower ribosome elongation rate modulated by low codon optimality affects the stability of mRNAs in yeast [12]. This led us to examine whether decelerated ribosomes could be observed especially in regions where GC3-content was low. Because we speculated that a single codon would be insufficient in eliciting any noticeable effects on the speed of the ribosome, we divided each CDS into 25 bins from start codon to stop codon and summed GC3-content within each bin. From the PCA, PC1 factor loadings of the codons were indicative of how much a particular codon contributed to the AT3-GC3 grouping i.e. instability-stability (**Fig EV2A**). Therefore as a measure of estimating ribosome occupancy, the factor loading scores of the codons from the first principal component were utilized to derive codon bias-derived occupancy scores. We then compared these scores with corresponding ribosome occupancies derived from ribosome profiling [26]. Ribosome occupancy obtained from HEK293 cells growing under physiological conditions generally coincided with codon bias-derived occupancy (Fig 2A). These measurements were highly reproducible between replicates of ribosome profiling experiments across the transcriptome (R^2^ = 0.750, 16,423 transcripts) (**Fig EV2B**). We observed a significantly better prediction of ribosome occupancy by codon bias-derived occupancy scores than that derived from scrambled codon bias-derived occupancy scores (Fig 2B). Indeed, representative transcripts showed a good correlation between our codon bias-derived occupancy scores and ribosome occupancy as exemplified by EIF2B2, DYNC1LI2, and IDH3G transcripts (**Fig EV2C**).

**Figure 2.**
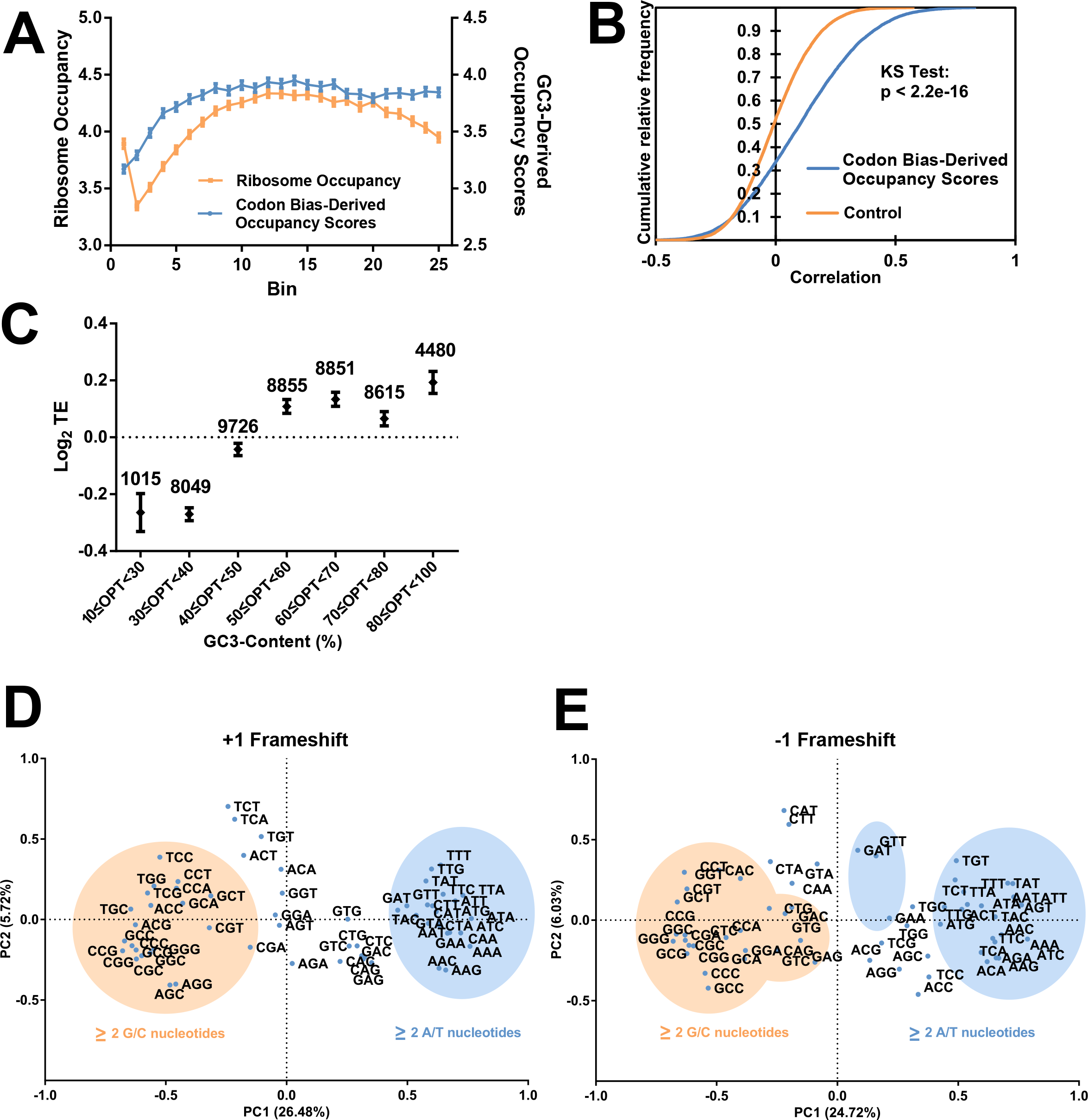
Ribosome profiling reveals that ribosome occupancy is moderately correlated with codon bias. **A.** Average ribosome occupancy and their respective codon bias-derived occupancy scores across the CDS of transcript (in 25 bins). Ribosome occupancy for 16,423 transcripts and their respective codon bias-derived occupancy were firstly binned into 25 bins and the mean occupancy was calculated for each bin. **B.** Cumulative distribution plots showing the distributions of correlations between ribosome occupancy and codon bias-derived occupancy. Correlations obtained from the ribosome occupancy and scrambled codon bias-derived occupancy served as the control. A Kolmogorov–Smirnov test was performed between the codon bias-derived occupancy and the control group. **C.** Comparison of average transcript translation efficiencies (TEs) across their respective GC3-content ranges. Number of transcripts within each gene optimality range is indicated above their respective points. **D, E.** Principal component analysis of CDS codon frequencies of protein-coding genes derived from a +1 frameshift (**D**) and a −1 frameshift (**E**). Shaded ellipses indicate codons which are GC-rich (orange) and AT-rich (blue). **Data information:** In (**A, C**), error bars represent the 95% confidence intervals.

Although translation elongation and initiation are distinct steps, previous literature has suggested that optimal codons are also enriched in mRNAs with high translation [27]. Ribosome footprint reads normalized by mRNA abundances from RNA-Seq enables the calculation of translation efficiency which in turn is also generally regarded as the translation initiation rate [28]. Therefore, to establish the link between translation status and codon bias, we calculated the translation efficiency (TE)—ribosome footprints normalized by mRNA abundance. Indeed, our results showed that mRNAs with high GC3-content generally possessed high TE (Fig 2C).

To verify if GC3 and AT3 codons were indeed associated with stability and instability respectively, we performed PCA on +1 and −1 frameshifted CDS sequences genome-wide and show that the GC3-AT3 demarcation was abolished (Fig 2D **and** E). Interestingly, we found that GC-rich (two or three G/C bases) and AT-rich (two or three A/T bases) codons contributed strongly to PC1 of the frameshifted data showing that GC/AT content is a natural consequence GC3-AT3 usage (Fig 1D **and Fig EV1C**). This phenomenon was also observed when we compared the frameshifted codon frequencies with mRNA decay rates from Fig 1C.

Thus far, we show that GC3 and AT3 codons are associated with mRNA stability, ribosome translation speed and efficiency therefore suggesting that the former and latter can be designated into optimal and non-optimal codons respectively.

### Codon bias affects mRNA stability

We then experimentally validated our bioinformatics observations of GC3 and AT3 codons in human cells. We developed a scheme based on the PC1 factor loadings in which we previously utilized in our ribosome profiling analysis (**Fig EV2A**). Based on this scheme, codons could be optimized and de-optimized with regard to GC3 content within their codon boxes *i.e.* synonymous substitutions (**Fig EV3A**). Single box codons such as TGG (Trp) and ATG (Met) would remain unchanged. We synthesized two independent genes (*REL* and *IL6*) with differential GC3-content (**Fig EV3B**) and examined the stability of these reporter RNA in HEK293 cells utilizing the Tet-off system (Fig 3A). As expected, the optimized transcripts of *REL* and *IL6* were more stable than their wild-type counterparts. Additionally, the decay rate of the de-optimized *IL6* reporter was faster, confirming that low GC3-content transcripts were unstable.

**Figure 3.**
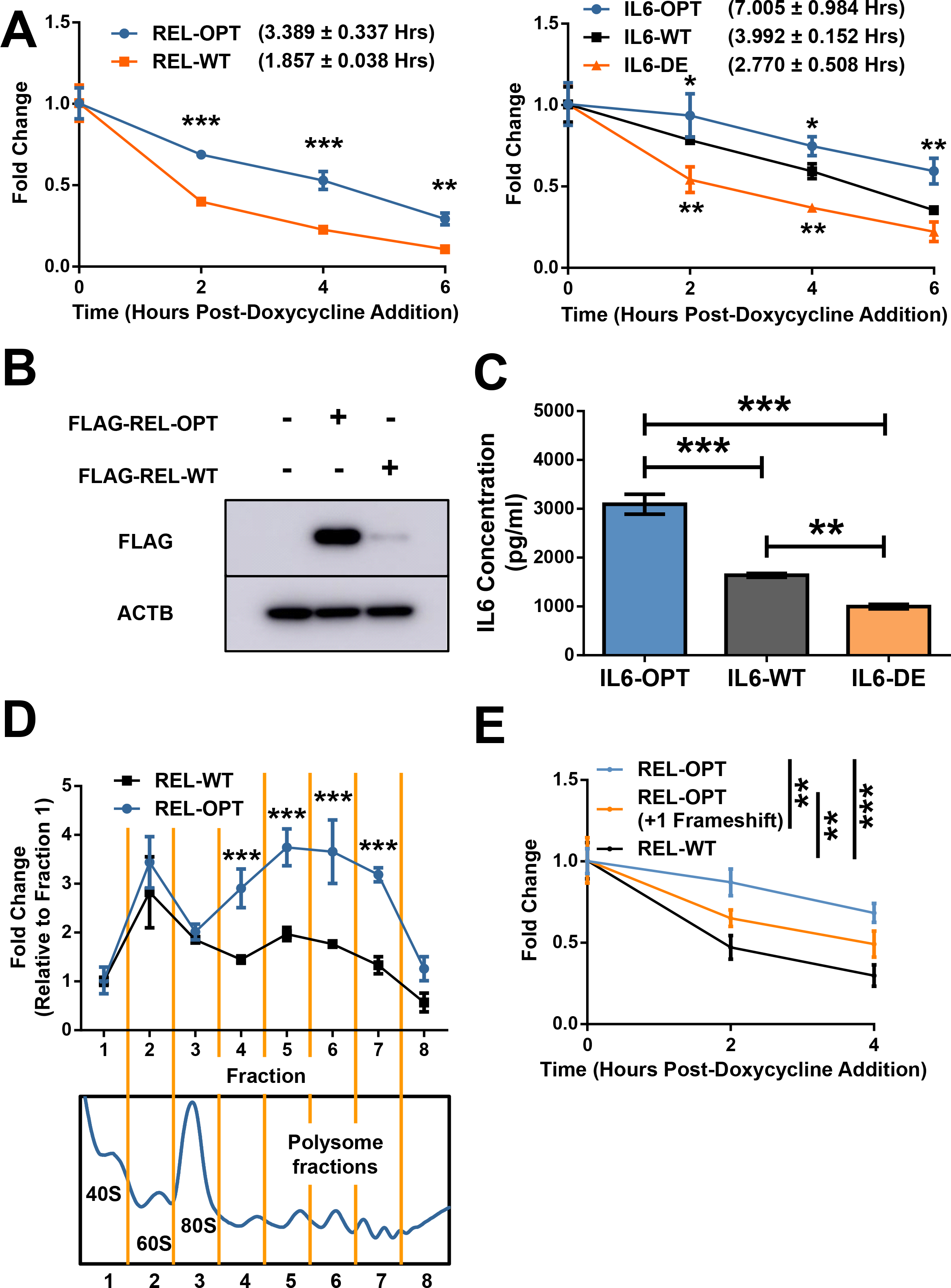
GC3-content of transcripts determine their fate. **A.** HEK293 Tet-off experiments showing the degradation of *REL-OPT* and *REL-WT* transcripts (left) as well as *IL6-OPT, IL6-WT* and *IL6-DE* transcripts (right), post-doxycycline addition. **B.** Representative immunoblot of FLAG-tagged REL-OPT and REL-WT in HEK293T cells transfected with either empty plasmids, plasmids bearing REL-OPT or REL-WT. The immunoblot is representative of 3 independent experiments. ACTB is shown as the loading controls. **C.** ELISA of secreted IL6 concentrations of IL6-OPT, IL6-WT and IL6-DE from HEK293T cells transfected with plasmids bearing IL6-OPT, IL6-WT and IL6-DE. **D.** Fold changes of REL-OPT and REL-WT transcript levels (top) relative to their abundances from fraction 1 as detected by qPCR across polysome fractions (below). Data represents the mean ± SD for 3 biological replicates. **E.** HEK293 Tet-off experiments showing the degradation of *REL-OPT, REL-OPT (+1 Frameshift)* and *REL-WT* transcripts post-doxycycline addition. **Data information:** In (**A, E**), data is representative of 3 independent experiments each with 3 replicates. The data represents the mean ± SD for 3 replicates. A two-way ANOVA with Holm-Sidak multiple comparisons was performed. P-values are denoted as follows: p < 0.05 (*), p<0.01 (**) and p<0.001 (***). The half-lives of the respective transcripts are indicated in brackets. In (**C**), the data is representative of 3 independent experiments each with 3 replicates. The data represents the mean ± SD for 3 replicates. A one-way ANOVA with Tukey’s multiple comparisons was performed between samples where, p<0.01 (**) and p<0.001 (***).

In addition to the RNA stability, higher GC3-content was also associated with higher translation efficiency (Fig 2C), thereby increasing protein production. Indeed, the protein abundance of the optimized *REL* reporter was higher than *REL-WT* even after normalization of protein abundance by mRNA levels (Fig 3B, **Fig EV3C**). Using enzyme-linked immunosorbent assay (ELISA), we observed that expression of *IL6-OPT* resulted in a 1.5-fold and 2-fold significantly higher level of IL6 compared to its WT and *IL6-DE*, respectively (Fig 3C). In a similar fashion, normalization of IL6 protein abundance by mRNA levels revealed that translation efficiency of the optimized *IL6* reporter was higher than its WT and de-optimized reporter counterparts (**Fig EV3D**). We tested our REL reporters in HeLa cells and show that the high protein abundance of *REL-OPT* could also be observed (**Fig EV3E**). Moreover, polysome fractionation and subsequent qPCR analysis revealed that within the polysome fractions, *REL-OPT* transcript amounts were significantly higher than *REL-WT* transcripts, suggesting that *REL-OPT* was translated more efficiently than *REL-WT* (Fig 3D). Thus far, our results validate the bioinformatics analyses and show that GC3 and AT3 codons can be designated as optimal and non-optimal codons.

We then synthesized a +1 frameshifted version of the REL-OPT transcript, removing any potential stop codons which would have resulted in premature termination of transcription and measured its stability via the Tet-off system (Fig 3E). This frameshifted version, while possessing a lower GC3-content, retained the same GC-content as its in-frame counterpart (*REL-OPT*) (**Fig EV3B**). Surprisingly, the frameshifted version, was still more stable than the WT form, yet less stable compared to its in-frame optimized counterpart, suggesting that GC-content could be an additional determinant of stability. Taken together, our results show that codon bias encompasses two modes of mRNA regulation, GC3- and GC-content dependent.

### RNA binding proteins differentially bind to transcripts of varying degrees of codon bias

Having shown that high optimality content inevitably accords high GC-content which in turn promotes mRNA stability, we wondered if there were RNA binding proteins (RBPs) which scrutinize, discriminate or even affect an mRNA’s fate. To identify RBPs which were either bound to transcripts bearing high or low optimality, we performed a ribonucleoprotein immunoprecipitation-based approach termed ISRIM (*In vitro* <u>S</u>pecificity based <u>R</u>NA Regulatory protein Identification <u>M</u>ethod) [29]. Lysates of HEK293 cells were mixed with FLAG peptide-conjugated *REL* and *IL6* transcripts of high and low optimality and their interacting proteins were determined using mass spectrometry. We then calculated the fold changes based on the abundance of RBPs bound to *REL-WT* with respect to *REL-OPT* (Fig 4A).

**Figure 4.**
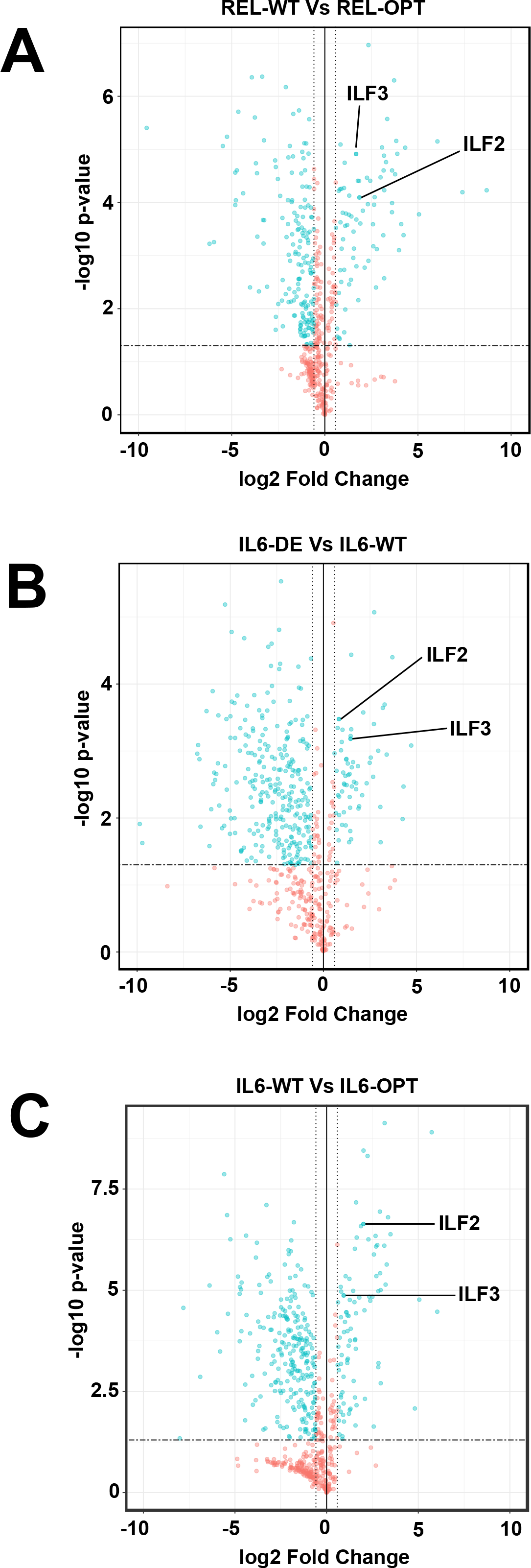
RNA binding proteins bind differentially to transcripts with different levels of GC3-content. **A, B, C.** Volcano plots showing the enrichment of RBPs which bind to *REL-WT* relative to *REL-OPT* transcripts (**A**), *IL6-DE* relative to *IL6-WT* transcripts (**B**) and *IL6-WT* relative to *IL6-OPT* transcripts (**C**). **Data information:** In (**A, B, C**), vertical dotted lines indicate a 1.5-fold enrichment while horizontal dashed lines indicate the p-value cut-off of 0.05. Points shaded in blue indicate RBPs which have a differential fold change of more than 1.5 and p<0.05.

As IL6 transcripts possessed three levels of GC3-content (OPT, WT, DE), we defined high GC3-content binding RBPs based on the RBP enrichment of *IL6-WT* to *IL6-DE* (Fig 4B) as well as *IL6-OPT* compared *to IL6-WT* (Fig 4C). Similarly, we defined low GC3-content binding RBPs based on the RBP enrichment of *IL6-DE* compared to *IL6-WT* (Fig4B) as well as *IL6-WT* to *IL6-OPT* (Fig 4C). By selecting common RBPs belonging to each group, we defined a set of RBPs which bound differentially to high GC3 and low GC3 IL6 transcripts respectively (**Fig EV4A**) We then selected RBP candidates which were specifically enriched with either low or high GC3 transcripts common to both REL and IL6 ISRIM experiments (**Fig EV4B**, **Dataset EV2**). In all, we show that RBPs can differentiate between transcripts of high GC3 and low GC3 content.

### ILF2 regulates the stability of low GC3 / high AT3 transcripts

We investigated the role of RBPs in modulating the stability of transcripts with different codon bias. Of interest were ILF2 and ILF3, RPBs identified from the list of RBPs interacting exclusively with low optimality transcripts. ILF2 and ILF3, also known as NF45 and NF90/NF110, respectively, are well known to function dominantly as heterodimers which bind double stranded RNA. ILF3 has been extensively studied, having shown to bind to AU-rich sequences in 3′ UTR of target RNA to repress its translation [30]. We hypothesize that the binding of ILF2 and ILF3 to their targets occur as low optimality transcripts are inadvertently AU-rich. Here we focused on the effects of these RBPs on low optimality transcripts. Firstly, using published RIP-seq data of ILF2 in two multiple myeloma cell lines, H929 and JJN3, we observed that ILF2 interacts with low optimality transcripts (**Fig EV5A**) [31; Data ref: Marchesini et al. 2017]. Additionally, we analysed RNA-Seq data obtained from the ENCODE project of K562 cells treated by CRISPR interference targeting ILF2 [32; Data ref: ENCODE Database ENCSR073QLQ]. Strikingly, we observed that transcripts that possessed low optimality scores were upregulated whereas transcripts that possessed high optimality scores were downregulated (Figure 5A, **Fig EV5B**). The abundance changes of representative mRNAs by ILF2 knockdown were antiparallel to their GC3-content (**Fig EV5C**).

**Figure 5.**
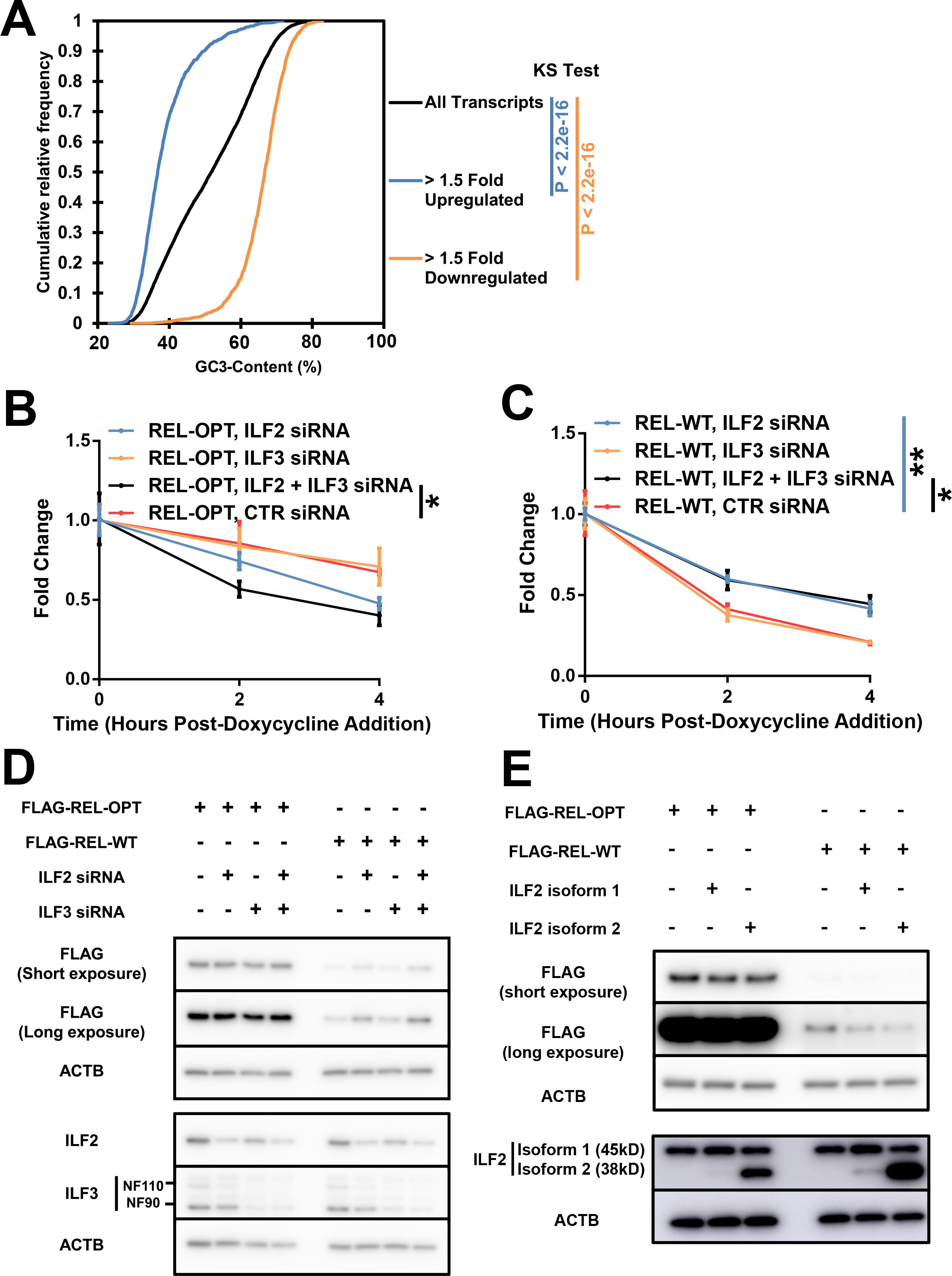
ILF2 regulates the stability of low GC3 / high AT3 transcripts. **A.** Cumulative distribution plots showing the difference in distribution of transcript optimality between upregulated and downregulated transcripts in K562 cells subject to ILF2 CRISPR interference targeting ILF2. **B, C.** HEK293 Tet-off experiments showing the degradation of *REL-OPT* (**B**) and *REL-WT* (**C**) transcripts with ILF2 and ILF3 siRNA and Control (CTR) siRNA treatment, post-doxycycline addition. **D.** Representative immunoblot of FLAG-tagged REL-OPT and REL-WT expressed in HEK293T cells under ILF2 and ILF3 siRNA treatment. The immunoblot is representative of 3 independent experiments. ACTB was shown as loading controls. **E.** Representative immunoblot of FLAG-tagged REL-OPT and REL-WT in HEK293T cells co-expressed with two different isoforms of ILF2. The immunoblot is representative of 3 independent experiments. ACTB was shown as loading controls. **Data information:** In (**A**), Kolmogorov–Smirnov tests were performed on the upregulated and downregulated groups against the control group. P-values are denoted (right). In (**B, C**), data is representative of 3 independent experiments in which the data represents the mean ± SD for 3 biological replicates. Unpaired t-tests were performed between samples where, p < 0.05 (*) and p<0.01 (**).

To confirm our observations, we examined the stability of FLAG-tagged versions of *REL-OPT* and *REL-WT* in the Tet-off system after ILF2 and ILF3 knockdown via siRNA (Fig 5B-C). Interestingly, we observed that the optimized reporter was more unstable under the knockdown of both ILF2 and ILF3 whereas the WT reporter was more stable with the knockdown of ILF2 only and a combination of both ILF2 and ILF3. No change in stability was observed with ILF3-only knockdown WT reporters suggesting that the increase in the stability of the WT reporter was due to the knockdown of ILF2, not ILF3. In agreement with this, we found a significant increase in protein levels of REL-WT when cells were treated with ILF2- and ILF3-targeting siRNA (Fig 5D **and Fig EV5D**). However, despite seeing a decrease in stability of the GC3-optimized reporter under both ILF2 and ILF3 knockdown, we were unable to observe this change at the protein level.

Focusing our attention on ILF2, we expressed FLAG-tagged versions of *REL-OPT* and *REL-WT*, along with the two isoforms of ILF2 and detected the reporter protein levels via western blot. A significant decrease in band intensity was observed for the REL-WT bands when both isoforms of ILF2 were expressed, whereas the amount of REL-OPT was not changed (Fig 5E **and Fig EV5E**). Taken together, our results suggest that ILF2 targets mRNA transcripts with low GC3-content (and inadvertently low GC-content) to induce their decay.

## Discussion

This study provides a framework describing codon bias-mediated RNA decay in humans. We first show that GC3 codons are associated with stability and AT3 codons with instability. We quantified codon bias by calculating the GC3 content within the CDS of genes and showed that GC3-content is strongly correlated with RNA stability and amount of protein expressed. In general, the use of optimal GC3 codons correlated with higher GC-content at a genome-wide level. We then show a modest agreement between codon bias-derived scores and ribosome occupancy as determined by ribosome profiling. Using GC3-optimized and de-optimized reporters we validate our bioinformatics observations *in vitro*. Screening of RNA binding proteins revealed that ILF2, which interacts with transcripts with low GC3-content, induces their degradation. Taken together, we conclude that gene expression can be shaped by codon bias and inevitably by GC/AU-content through the modulation of mRNA stability in human cells.

### Investigating the System of Codon Bias in Humans

Since translation elongation is affected by tRNA availability, the tRNA adaptation index (tAI), which is based on genomic tRNA copy number, has been used as a surrogate for codon optimality. However, in contrast to yeast, tRNA copy number in genome is not always correlated with tRNA abundance in higher eukaryotes [33]. Hence, this metric is less suitable for quantifying codon optimality in humans.

Independent of tRNA-based metrics, we addressed these challenges by utilizing an unsupervised learning algorithm, PCA, to identify features in that were mRNA-intrinsic. In the PCA of both yeast and humans, we demonstrated that the first principal component mirrored optimal/non-optimal assignments. We also show that the codon bias is different between these two organisms (Fig 1B, **Fig EV1A**). In humans, the classification of codons into AT3 and GC3 groups was striking, but the percentage by which it accounts for its variation however was modest. From the PCA, the first and second principal components only explain a quarter of total variance in codon frequencies (Fig 1B), implying that other factors that explain bias of codon frequency possibly remains in human cells.

Assuming that evolution drives the selection of codons, synonymous codon usage in different organisms must be fine-tuned over time to achieve precise expression levels of mRNA and eventually proteins in essential physiological process. It is therefore plausible that high GC3/AT3-content or GC/AT-content in mRNA is selected for to modulate transcript stability in essential physiological processes, but subject to constraints by amino sequence. Indeed, we show that transcripts with high and low GC3-content were linked to particular physiological and cellular processes (**Fig EV1E and F**). In a particular study, Gingold and colleagues argue that tRNA abundances vary in proliferating and differentiating cell types [34]. Interestingly, they showed that codons preferred by cell cycling genes were AT3 codons while pattern-specification preferred codons tended to be GC3 codons—in agreement with our GO analyses. In *Drosophila*, the correlation between codon optimality and mRNA stability has been demonstrated to be attenuated in neural development, possibly allowing the effect of trans-acting factors to dominate development [22]. c-Rel, a protein encoded by the *REL* gene and a canonical nuclear factor κB (NF-κB) subunit, is expressed abundantly in differentiated lymphoid cells and has been shown to be vital in thymic regulatory T cell development in addition to controlling cancer via activated regulatory T cells [35,36]. Given the inherent low optimality and associated instability of *REL* in its WT form (Fig 3A), we wonder if besides transcriptional control of *REL*, could there be other post-transcriptional regulation systems at play. Further studies would be necessary to investigate if codon optimality or codon optimality-associated RBPs modulate *REL* gene expression.

Our results show that codon bias affects ribosome occupancy to a significant extent (Fig 2B). Taking into account our *in vitro* experiment results together with the ribosomal profiling results, we suggest that GC3 and AT3 codons are synonymous with optimal and non-optimal codons. Additionally, our study along with others’ suggests that slower elongation of ribosome is a key feature of mRNA stability. Although codon optimality is a dominant factor in general, other factors may also be involved in decelerated ribosomes, such as secondary structures [37,38]. These obstacles for ribosome elongation are reversible and dynamically regulated by RNA helicases [39,40]. Further studies will be required to elucidate the role of RNA secondary structures and helicases and their relevance to codon bias and mRNA stability decay.

### The stability of mRNA can be modulated by RBPs which bind AU-rich sequences

Whereas AU-rich elements (AREs) in the 3′ UTR have been traditionally targeted by RBPs, we found that coding regions are also targeted by ARE-recognizing RBPs. The identification of the heterodimeric complex consisting of ILF2 and ILF3 among others shows that a wide array of RBPs recognizes low optimality (AU-rich) sequences (Fig 4). Here, we show that ILF2 destabilizes AU-rich, GC-poor transcripts. At the protein level however, knockdown ILF2 and ILF3 show better stabilization of GC3-poor transcripts even when compared to ILF2 knockdown alone. Interestingly, Kuwano and colleagues show that NF90, the shorter isoform of ILF3, specifically targets AU-rich sequences in mRNA 3’UTRs and represses their translation, not stability [30]. Taking into consideration reports that ILF2 and ILF3 can function independently of each other [41–43], it is possible that ILF2 and ILF3 regulate the fate of mRNA differently. Our screens also detected HNRNPD/AUF1, which destabilizes transcripts via recognition of AU-rich motifs [44], binding to low optimality mRNAs (**Dataset EV2**). These observations emphasize the importance of AU-content, which is strongly connected with low optimality, in RNA destabilization. However, we could not exclude the possibility that these factors induced the degradation of AU-rich transcripts independent of the model proposed by Presnyak and Radhakrishnan [12,13] as our RBP identification method was not fully reflective of the active translational status required for co-translational degradation of mRNA transcripts. Further studies would be necessary to discern if these or other factors act as sensors of codon optimality during translation.

In conclusion, in human cells, the redundancy of the genetic code allows the choice between alternative codons for the same amino acid which may exert dramatic effects on the process of translation and mRNA stability. In our experiments, we show that two modes of mRNA regulation exist – GC3 and GC-content dependent. This system potentially confers freedom for calibrating protein and mRNA abundances without altering protein sequence. Beginning from our exploratory analysis, we have developed an approach to quantify codon bias and demonstrate that beneath the redundancy of codons, exists a system which modulates mRNA and consequently, protein abundance.

## Materials and Methods

### Cell Cultures, Growth, and Transfection Conditions

HEK293T cells were maintained in Dulbecco’s modified eagle medium (DMEM) (Nacalai Tesque), supplemented with 10% (v/v) fetal bovine serum. HEK293 Tet-off cells were maintained in Minimum Essential Medium Eagle - Alpha Modification (α-MEM) (Nacalai Tesque), supplemented with 10% (v/v) Tet-system approved fetal bovine serum (Takara Bio) and 100 µg/ml of G418 (Nacalai Tesque). For REL and IL6 overexpression experiments, plasmids were transfected using PEI MAX (Polysciences Inc). For co-transfection of ILF2 siRNA with REL plasmids, Lipofectamine 2000 was used as per manufacturer’s protocol. ILF2 siRNA which targeted ILF2 at exons 8 and 9 were Silencer Select siRNA, S7399 (Ambion, Life Technologies).

### Plasmid Construction

Codon optimized-*REL* (*REL-OPT*), *IL6* (*IL6-OPT*) and codon de-optimized *IL6* (*IL6-DE*) sequences were synthesized as gBlocks Gene Fragments (Integrated DNA Technologies) (**Dataset EV3**). The *REL-OPT (+1 Frameshift)* sequence was constructed by adding a +1 frameshift just after the start codon. Resulting stop codons were removed to ensure no premature termination. These sequences and corresponding WT sequences were polymerase chain reaction (PCR) amplified (with the inclusion of a FLAG tag for *REL* sequences) and inserted into the pcDNA3.1(+) vector (Invitrogen) and pTRE-TIGHT vector (Takara Bio). The sequences were confirmed via restriction enzyme digest and sequencing.

### Tet-Off Assay

HEK293 Tet-off cells (Clontech) were transfected with pTRE-TIGHT plasmids bearing the (de)optimized and WT sequences and incubated overnight at 37ºC. Transcriptional shut-off for the indicated plasmids was achieved by the addition of doxycycline (LKT Laboratories Inc.) to a final concentration of 1 µg/ml. Samples were harvested at the indicated timepoints after the addition of doxycycline.

### RNA Extraction, Reverse Transcription PCR, and Quantitative Real-time PCR

Total RNA was isolated from cells using TRIzol reagent (Invitrogen) as per manufacturer’s instructions. Reverse transcription was performed using the ReverTra Ace qPCR RT Master Mix with gDNA remover kit (Toyobo) as per manufacturer’s instructions. cDNA was amplified with PowerUp SYBR Green Master Mix (Applied Biosystems) and quantitative real-time PCR (qPCR) was performed on the StepOne Real-Time PCR System (Applied Biosystems). Human GAPDH abundance was used for normalization. The list of qPCR primers can be found in **Dataset EV3**.

### Sucrose Gradient Centrifugation (Polysome Profiling)

HEK293T were transfected with equal concentrations of *REL-OPT* and *REL-WT* plasmids. Cells were lysed the next day in polysome buffer [20 mM 4-(2-hydroxyethyl)-1-piperazineethanesulfonic acid (HEPES-KOH) (pH 7.5], 100 mM KCl, 5 mM MgCl_2_, 0.25% (v/v) Nonidet P-40, 10 µg/ml cycloheximide, 100 units/ml RNase inhibitor, and protease inhibitor cocktail (Roche)]. Lysates were loaded on top of a linear 15%–60% sucrose gradient [15%–60% sucrose, 20 mM HEPES-KOH [pH 7.5], 100 mM KCl, 5 mM MgCl_2_, 10 µg/ml cycloheximide, 100 units/ml RNase inhibitor, and protease inhibitor cocktail (Roche)]. After ultracentrifugation at 38,000 rpm for 2.5 h at 4ºC in a HITACHI P40ST rotor, fractions were collected from the top of the gradient and subjected to UV-densitometric analysis. The absorbance profiles of the gradients were determined at 254 nm. For disassociation of ribosome and polysome, EDTA was added to Mg^2+^-free polysome buffer and 15%–60% sucrose gradient at concentrations of 50 mM and 20 mM, respectively. For RNA analysis, RNA from each fraction was extracted via the High Pure RNA Isolation Kit (Roche) and subject to reverse transcription and qPCR.

### Immunoblot Analysis

Samples were lysed in RIPA buffer (20 mM Tris-HCl [pH 8], 150 mM NaCl, 10 mM EDTA, 1% Nonidet-P40, 0.1% SDS, 1% sodium deoxycholate, and cOmplete Mini EDTA-free Protease Inhibitor Cocktail [Roche]). Protein concentration was determined by the BCA Protein Assay (Thermo Fisher). Whole cell lysates were resolved by SDS-PAGE and transferred onto PVDF membranes (Bio-Rad). The following antibodies were used for immunoblot analysis: mouse monoclonal anti-FLAG (F3165, Sigma), mouse monoclonal anti-ILF2 (sc-365283, Santa Cruz Biotechnology), mouse anti-β-actin (sc-47778, Santa Cruz), and mouse IgG HRP linked F(ab’)_2_ fragment (NA9310, GE Healthcare). Luminescence was detected with a luminescent image analyser (Amersham Imager 600; GE Healthcare).

### ELISA

HEK293T cells were transfected with pcDNA3.1(+) plasmids bearing the (de)optimized and WT sequences and incubated overnight at 37ºC. Cell supernatant was aspirated and the cell monolayer washed with 1x PBS (pre-warmed at 37ºC). Pre-warmed DMEM was added to the monolayer and the cells incubated for 2 hr at 37ºC. Thereafter, the cell supernatant was harvested and centrifuged at 300 × g to pellet residual cells. The resulting supernatant was decanted and the concentration of secreted IL6 was measured by the human IL6 ELISA kit (Invitrogen) according to the manufacturer’s instructions.

### ISRIM (*In vitro* Specificity based RNA Regulatory protein Identification Method)

#### Preparation of bait RNAs

T7-tagged cDNA template was PCR amplified and subjected to *in vitro* transcription using a MEGAscript T7 kit (Applied Biosystems). Amplified cRNA was purified with an RNeasy Mini Kit (Qiagen) and then subjected to FLAG conjugation as described (10) with some modifications. Briefly, 60 μl of freshly prepared 0.1 M NaIO_4_ was added to 60 μl of 250 pmol cRNA, and the mixture was incubated at 0ºC for 10 min. The 3′ dialdehyde RNA was precipitated with 1 ml of 2% LiClO_4_ in acetone followed by washing with 1 ml acetone. The pellet was dissolved in 10 μl of 0.1 M sodium acetate, pH 5.2 and then mixed with 12 μl of 30 mM hydrazide–FLAG peptide. The reaction solution was mixed at room temperature for 30 min. The resulting imine-moiety of the cRNA was reduced by adding 12 μl of 1 M NaCNBH_3_, and then incubated at room temperature for 30 min. The RNA was purified with an RNeasy Mini Kit (Qiagen).

#### Purification and analysis of RNA-binding proteins

Purification and analysis of RNA-binding protein (RBP) were carried out as described [29] with some modifications. Briefly, HEK293T cells were lysed with lysis buffer [10 mM HEPES (pH 7.5), 150 mM NaCl, 50 mM NaF, 1 mM Na_3_VO_4_, 5 μg/ml leupeptin, 5 μg ml aprotinin, 3 μg/ml pepstatin A, 1 mM phenylmethylsulfonyl fluoride (PMSF), and 1 mg/ml digitonin] and cleared by centrifugation. The cleared lysate was incubated with indicated amounts of FLAG-tagged bait RNA, antisense oligos and FLAG-M2-conjugated agarose for 1 hr. The agarose resin was then washed three times with wash buffer [10 mM HEPES (pH 7.5), 150 mM NaCl, and 0.1% Triton X-100] and co-immunoprecipitated RNA and proteins were eluted with FLAG elution buffer [0.5 mg/ml FLAG peptide, 10 mM HEPES (pH 7.5), 150 mM NaCl, and 0.05% Triton X-100]. The bait RNA associated proteins were digested with lysyl endopeptidase and trypsin. Digested peptide mixture was applied to a Mightysil-PR-18 (Kanto Chemical) frit-less column (45 3 0.150 mm ID) and separated using a 0–40% gradient of acetonitrile containing 0.1% formic acid for 80 min at a flow rate of 100 nl/min. Eluted peptides were sprayed directly into a mass spectrometer (Triple TOF 5600+; AB Sciex). MS and MS/MS spectra were obtained using the information-dependent mode. Up to 25 precursor ions above an intensity threshold of 50 counts/s were selected for MS/MS analyses from each survey scan. All MS/MS spectra were searched against protein sequences of RefSeq (NCBI) human protein database using the Protein Pilot software package (AB Sciex) and its decoy sequences then selected the peptides FDR was <1%. Ion intensity of peptide peaks ware obtained using Progenesis QI for proteomics software (version 3 Nonlinear Dynamics, UK) according to the manufacturer’s instructions.

### Ribosome profiling and RNA-Seq

Ribosome profiling was performed according to the method previously described with following modifications [26]. RNA concentration of naïve HEK293T lysate was measured by Qubit RNA BR Assay Kit (Thermo Fisher Scientific). The lysate containing 10 µg RNA was treated with 20 U of RNase I (Lucigen) for 45 min at 25ºC. After ribosomes were recovered by ultracentrifugation, RNA fragments corresponding to 26-34 nt were excised from footprint fragment purification gel. Library length distribution was checked using a microchip electrophoresis system (MultiNA, MCE-202, Shimadzu).

For RNA-seq, total RNA was extracted from the lysate using TRIzol LS reagent (Thermo Fisher Scientific) and Direct-zol RNA Kit (Zymo research). Ribosomal RNA was depleted using the Ribo-Zero Gold rRNA Removal Kit (Human/Mouse/Rat) (Illumina) and the RNA-seq library was prepared using TruSeq Stranded mRNA Library Prep Kit (Illumina) according to the manufacturer’s instructions.

The libraries were sequenced on a HiSeq 4000 (Illumina) with a single-end 50 bp sequencing run. Reads were aligned to human hg38 genome as described [26,45]. The offsets of A-site from the 5′ end of ribosome footprints were determined empirically as 15 for 25-30 nt, 16 for 31-32 nt, and 17 for 33 nt. For RNA-seq, offsets were set to 15 for all mRNA fragments. For calculation of the ribosome occupancies, mRNAs with lower than one footprint per codon were excluded. For calculation of the translation efficiencies (TEs), we counted the number of reads within each CDS, and ribosome profiling counts were normalized by RNA-seq counts using the DESeq package [46]. Reads corresponding to the first and last five codons of each CDS were omitted from the analysis of TEs. The Custom R scripts will be available upon requests.

### Bioinformatics and Computational Analyses

#### Principal component analysis

CDS data (Human genes, GRCh38p12) was obtained from the Ensembl Biomart Database and the average human codon frequencies per transcript were calculated for all transcripts. The resulting data was subject to PCA analysis using the Python 3.4 environment via the factoextra program [47]. Finally, the data was trimmed to remove truncated sequences as well as sequences with non-canonical start codons to a final of 9898 genes.

#### Hierarchical clustering analysis

mRNA transcripts ranked in order of their half-lives, divided equally into 4 groups and their average half-lives within each group was calculated. The corresponding codon frequencies of transcripts within each group were averaged. Hierarchical clustering was performed using the average linkage method to cluster the codon frequencies in R using the ggplot2 program [48].

#### Quantification of GC3-content

To quantify GC3-content, we summed up the codon frequencies of GC3 codons and expressed the frequencies on a percentage scale.

#### Binning of ribosomal occupancy frequencies and calculation of codon bias-derived occupancy scores

To quantify codon bias for ribosome profiling, the factor loading scores of the codons from the first principal component were normalized linearly on a percentage scale from 0 to 1 where 0 corresponded to the codon with the lowest score (AAT) and 1 for the codon with the highest score (GCC) (**Fig EV2A**). Binning of the ribosome occupancies were performed in the R environment via a custom script. To calculate the corresponding codon bias-derived occupancy scores, we substituted the codon sequences of mRNA transcripts with their respective codon scores and in a similar fashion, binned the data into 25 bins. As the scores of codons should inversely reflect the ribosome occupancy (i.e. higher ribosome occupancy associated with lower codon scores), we calculated the reciprocal of the binned codon scores within each bin for all 25 bins to derive the codon bias-derived occupancy scores. Both ribosome occupancy and codon bias-derived occupancy scores were normalized on a linear scale and a Pearson correlation performed on each transcript. To exclude the possibility that the correlations were due to chance, we shuffled the bins for the codon bias-derived occupancy scores within each individual transcripts and calculated the Pearson correlation between shuffled and ribosomal occupancy data.

## Supporting information

Expanded View Figures 1-5

## Acknowledgements

The authors express their gratitude to all members of the laboratory of Medical Chemistry, Kyoto University, for their kind advice and discussions. DNA libraries were sequenced by the Vincent J. Coates Genomics Sequencing Laboratory at UC Berkeley, supported by NIH S10 OD018174 Instrumentation Grant. Computations were supported by Manabu Ishii, Itoshi Nikaido, and the Bioinformatics Analysis Environment Service on RIKEN Cloud at RIKEN ACCC.

This work was supported by the JSPS KAKENHI (18H05278), AMED-CREST from Japan Agency for Medical Research and Development and the JSPS through Core-to-Core Program.

This work was supported by Joint Usage/Research Center program of Institute for Frontier Life and Medical Sciences, Takeda Science Foundation, the Uehara Memorial Foundation.

S.I. was supported by Grant-in-Aid for Scientific Research on Innovative Areas “nascent chain biology” (JP17H05679) and Grant-in-Aid for Young Scientists (A) (JP17H04998) from JSPS, the Pioneering Projects ("Cellular Evolution") and the Aging Project from RIKEN, and Takeda Science Foundation.

## Author contributions

FH wrote the manuscript; together with SFY performed the experiments and analyzed the data. MY provided insightful comments and proofreading for the manuscript. YM and ML performed the mRNA decay experiments. YS, SI performed the ribosomal profiling and proofreading of the manuscript.SA and TN performed the ISRIM experiments. AV provided advice and bioinformatics expertise. AF and TF performed the polysome profiling experiments. OT supervised and designed the experiments.

**Expanded View Figure 1**

**A.** Principal component analysis of the CDS codon frequencies of protein-coding genes in S. cerevisiae. PC1 and PC2 indicate the first and second principal components.

**B.** F1 factor loadings of codons from the yeast dataset ranked from the highest to the lowest. The optimal and non-optimal designation at the bottom of the figure refers to the designation according to Presnyak and colleagues [12].

**C.** Pearson correlation between GC-content and GC3-content for 9666 protein-coding genes.

**D.** Violin plots visualizing the distribution of mRNA half-lives across their respective GC3-content brackets.

**E.** Gene ontology analysis (biological processes) of the top 5% ranked genes in terms of gene GC3-content

**F.** Gene ontology analysis (biological processes) of the bottom 5% ranked genes in terms of gene GC3-content

**Data information:** In (**D**), the box plots within each figure are indicative of the median and interquartile ranges.

**Expanded View Figure 2**

**A.** PC1 factor loadings of codons from the human dataset ranked from the highest to the lowest (bottom) and their corresponding normalized factor loadings after linear normalization onto a percentage scale (top).

**B.** Pearson correlation between the correlations of derived from comparison of ribosome occupancy and codon bias-derived scores for two ribosome profiling sample replicates.

**C.** Three example transcripts (EIF2B2, DYNC1LI2 and IDH3G) which demonstrate high correlation between ribosome occupancy and codon bias-derived scores (left) as well as their corresponding Pearson correlations over 25 bins (right).

**D, E.** Hierachical clustering analysis of half-lives of mRNA and their CDS codon frequencies after a +1 frameshift (**D**) and −1 frameshift (**E**). The transcripts were ranked according to their half-lives and divided equally into quartiles. The respective codon frequencies of each group were then averaged. Codon highlights indicate codons which are GC-rich (yellow) and AT-rich (blue).

**Expanded View Figure 3**

**A.** Example of how transcript GC3-optimization and deoptimization was performed to generate GC3-optimized and de-optimized versions of *REL* and *IL6* transcripts.

**B.** GC3- and GC-content of *REL-OPT/WT*, *REL-OPT (+1 Frameshift)*, and *IL6-OPT/WT/DE* transcripts.

**C.** Protein abundance of immunoblot of FLAG-tagged REL-OPT and REL-WT in HEK293T cells transfected with either empty plasmids, plasmids bearing REL-OPT or REL-WT (corresponding to Fig 3B). The protein abundance was normalized by respective mRNA levels.

**D.** Protein abundance as determined by ELISA of IL6-OPT, IL6-WT and IL6-DE in HEK293T cells transfected with either empty plasmids, plasmids bearing IL6-OPT, IL6-WT or IL6-DE (corresponding to Fig 3D). The protein abundance was normalized by respective mRNA levels. The ELISA quantification is representative of 3 independent experiments.

**E.** Representative immunoblot of FLAG-tagged REL-OPT and REL-WT expressed in HeLa cells (left). The immunoblot is representative of 3 independent experiments.

**Data information:** In (**C**), the densitometry data is representative of 3 independent experiments. Unpaired t-tests were performed within the REL-OPT and REL-WT samples, p<0.05 (*). In (**D**), a one-way ANOVA with Tukey’s multiple comparisons was performed between samples where, p<0.01 (**) and p<0.001 (***).

**Expanded View Figure 4**

**A.** Venn diagram indicating the number of RBPs identified from the IL6 ISRIM experiments.

**B.** Venn diagram indicating the number of RBPs identified from the REL and IL6 ISRIM experiments.

**Expanded View Figure 5**

**A.** Cumulative distribution plots showing the GC3-content distribution of transcripts bound to by ILF2 in H929 (top) and JJN3 cells (bottom). Kolmogorov–Smirnov tests were performed on the ILF2 RIP group against the control group.

**B.** Scatterplot of the RPKM values of mRNA transcripts in K562 cells subject to ILF2 CRISPR interference and its corresponding WT control. mRNA transcripts were colored according to their GC3-content.

**C.** Fold changes of example mRNA representing low, average and high GC3-content transcripts from the RPKM values of mRNA transcripts in K562 cells subject to ILF2 CRISPR interference.

**D.** Densitometric analysis of immunoblot of FLAG-tagged REL-OPT and REL-WT expressed in HEK293T cells under ILF2 and ILF3 siRNA treatment (corresponding to Fig 5D).

**E.** Densitometric analysis of immunoblot of FLAG-tagged REL-OPT and REL-WT expressed in HEK293T cells co-expressed with two different isoforms of ILF2 (corresponding to Figure 5E)

**Data information:** In (**C**), data represents the mean ± SD for 2 biological replicates. In (**D, E**), densitometry data is representative of 3 independent experiments. A one-way ANOVA with Tukey’s multiple comparisons was performed within the REL-OPT and REL-WT samples. P-values are denoted as follows, p < 0.05 (*), p<0.01 (**) and p<0.001 (***).

**Dataset EV1.** List of genes and their corresponding GC3- and GC-content.

**Dataset EV2.** List of RBPs identified in ISRIM experiments.

**Dataset EV3.** List of qPCR primers and their corresponding sequences.

## References

1. Huang L, Lou C-H, Chan W, Shum EY, Shao A, Stone E, Karam R, Song H-W, Wilkinson MF (2011) RNA Homeostasis Governed by Cell Type-Specific and Branched Feedback Loops Acting on NMD. Molecular Cell 43: 950–961.

2. Mino T, Murakawa Y, Fukao A, Vandenbon A, Wessels H-H, Ori D, Uehata T, Tartey S, Akira S, Suzuki Y, et al. (2015) Regnase-1 and Roquin Regulate a Common Element in Inflammatory mRNAs by Spatiotemporally Distinct Mechanisms. Cell 161: 1058–1073.

3. Yoshinaga M, Nakatsuka Y, Vandenbon A, Ori D, Uehata T, Tsujimura T, Suzuki Y, Mino T, Takeuchi O (2017) Regnase-1 Maintains Iron Homeostasis via the Degradation of Transferrin Receptor 1 and Prolyl-Hydroxylase-Domain-Containing Protein 3 mRNAs. Cell Rep 19: 1614–1630.

4. Leppek K, Das R, Barna M (2018) Functional 5′ UTR mRNA structures in eukaryotic translation regulation and how to find them. Nat Rev Mol Cell Biol 19: 158–174.

5. Cheng J, Maier KC, Avsec Ž, Rus P, Gagneur J (2017) Cis-regulatory elements explain most of the mRNA stability variation across genes in yeast. RNA 23: 1648–1659.

6. Vogel C, Marcotte EM (2012) Insights into the regulation of protein abundance from proteomic and transcriptomic analyses. Nat Rev Genet 13: 227–232.

7. Zhou T, Weems M, Wilke CO (2009) Translationally Optimal Codons Associate with Structurally Sensitive Sites in Proteins. Mol Biol Evol 26: 1571–1580.

8. Sharp PM, Li WH (1987) The codon Adaptation Index--a measure of directional synonymous codon usage bias, and its potential applications. Nucleic Acids Res 15: 1281–1295.

9. Reis M dos, Savva R, Wernisch L (2004) Solving the riddle of codon usage preferences: a test for translational selection. Nucleic Acids Res 32: 5036–5044.

10. dos Reis M, Wernisch L, Savva R (2003) Unexpected correlations between gene expression and codon usage bias from microarray data for the whole Escherichia coli K-12 genome. Nucleic Acids Res 31: 6976–6985.

11. Pechmann S, Frydman J (2013) Evolutionary conservation of codon optimality reveals hidden signatures of co-translational folding. Nat Struct Mol Biol 20: 237–243.

12. Presnyak V, Alhusaini N, Chen Y-H, Martin S, Morris N, Kline N, Olson S, Weinberg D, Baker KE, Graveley BR, et al. (2015) Codon optimality is a major determinant of mRNA stability. Cell 160: 1111–1124.

13. Radhakrishnan A, Chen Y-H, Martin S, Alhusaini N, Green R, Coller J (2016) The DEAD-Box Protein Dhh1p Couples mRNA Decay and Translation by Monitoring Codon Optimality. Cell 167: 122–132.e9.

14. Sweet T, Kovalak C, Coller J (2012) The DEAD-box protein Dhh1 promotes decapping by slowing ribosome movement. PLoS Biol 10: e1001342.

15. Dana A, Tuller T (2014) The effect of tRNA levels on decoding times of mRNA codons. Nucleic Acids Res 42: 9171–9181.

16. Gardin J, Yeasmin R, Yurovsky A, Cai Y, Skiena S, Futcher B Measurement of average decoding rates of the 61 sense codons in vivo. eLife 3:.

17. Drummond DA, Wilke CO (2008) Mistranslation-induced protein misfolding as a dominant constraint on coding-sequence evolution. Cell 134: 341–352.

18. Akashi H (1994) Synonymous Codon Usage in Drosophila Melanogaster: Natural Selection and Translational Accuracy. Genetics 136: 927–935.

19. Harigaya Y, Parker R (2016) Analysis of the association between codon optimality and mRNA stability in Schizosaccharomyces pombe. BMC Genomics 17: 895.

20. Lee Y, Zhou T, Tartaglia GG, Vendruscolo M, Wilke CO (2010) Translationally optimal codons associate with aggregation-prone sites in proteins. Proteomics 10: 4163–4171.

21. Mishima Y, Tomari Y (2016) Codon Usage and 3′ UTR Length Determine Maternal mRNA Stability in Zebrafish. Molecular Cell 61: 874–885.

22. Burow DA, Martin S, Quail JF, Alhusaini N, Coller J, Cleary MD (2018) Attenuated Codon Optimality Contributes to Neural-Specific mRNA Decay in Drosophila. Cell Rep 24: 1704–1712.

23. Boël G, Letso R, Neely H, Price WN, Wong K-H, Su M, Luff J, Valecha M, Everett JK, Acton TB, et al. (2016) Codon influence on protein expression in E. coli correlates with mRNA levels. Nature 529: 358–363.

24. Zerbino DR, Achuthan P, Akanni W, Amode MR, Barrell D, Bhai J, Billis K, Cummins C, Gall A, Girón CG, et al. (2018) Ensembl 2018. Nucleic Acids Res 46: D754–D761.

25. Murakawa Y, Hinz M, Mothes J, Schuetz A, Uhl M, Wyler E, Yasuda T, Mastrobuoni G, Friedel CC, Dölken L, et al. (2015) Gene Expression Omnibus GSE69153 (https://www.ncbi.nlm.nih.gov/geo/query/acc.cgi?acc=GSE69153)

26. McGlincy NJ, Ingolia NT (2017) Transcriptome-wide measurement of translation by ribosome profiling. Methods 126: 112–129.

27. Tuller T, Kupiec M, Ruppin E (2007) Determinants of Protein Abundance and Translation Efficiency in S. cerevisiae. PLOS Computational Biology 3: e248.

28. Iwasaki S, Ingolia NT (2016) Seeing translation. Science 352: 1391–1392.

29. Adachi S, Homoto M, Tanaka R, Hioki Y, Murakami H, Suga H, Matsumoto M, Nakayama KI, Hatta T, Iemura S, et al. (2014) ZFP36L1 and ZFP36L2 control LDLR mRNA stability via the ERK–RSK pathway. Nucleic Acids Res 42: 10037–10049.

30. Kuwano Y, Pullmann R, Marasa BS, Abdelmohsen K, Lee EK, Yang X, Martindale JL, Zhan M, Gorospe M (2010) NF90 selectively represses the translation of target mRNAs bearing an AU-rich signature motif. Nucleic Acids Res 38: 225–238.

31. Marchesini M, Ogoti Y, Fiorini E, Aktas Samur A, Nezi L, D’Anca M, Storti P, Samur MK, Ganan-Gomez I, Fulciniti MT, et al. (2017) Gene Expression Omnibus GSE83665 (https://www.ncbi.nlm.nih.gov/geo/query/acc.cgi?acc=GSE83665)

32. ENCODE Project Consortium (2012) (https://www.encodeproject.org/experiments/ENCSR073QLQ/)

33. Zheng G, Qin Y, Clark WC, Dai Q, Yi C, He C, Lambowitz AM, Pan T (2015) Efficient and quantitative high-throughput transfer RNA sequencing. Nat Methods 12: 835–837.

34. Gingold H, Tehler D, Christoffersen NR, Nielsen MM, Asmar F, Kooistra SM, Christophersen NS, Christensen LL, Borre M, Sørensen KD, et al. (2014) A Dual Program for Translation Regulation in Cellular Proliferation and Differentiation. Cell 158: 1281–1292.

35. Grinberg-Bleyer Y, Oh H, Desrichard A, Bhatt DM, Caron R, Chan TA, Schmid RM, Hayden MS, Klein U, Ghosh S (2017) NF-κB c-Rel Is Crucial for the Regulatory T Cell Immune Checkpoint in Cancer. Cell 170: 1096–1108.e13.

36. Oh H, Grinberg-Bleyer Y, Liao W, Maloney D, Wang P, Wu Z, Wang J, Bhatt DM, Heise N, Schmid RM, et al. (2017) An NF-κB Transcription-Factor-Dependent Lineage-Specific Transcriptional Program Promotes Regulatory T Cell Identity and Function. Immunity 47: 450–465.e5.

37. Pop C, Rouskin S, Ingolia NT, Han L, Phizicky EM, Weissman JS, Koller D (2014) Causal signals between codon bias, mRNA structure, and the efficiency of translation and elongation. Mol Syst Biol 10: 770.

38. Endoh T, Sugimoto N (2016) Mechanical insights into ribosomal progression overcoming RNA G-quadruplex from periodical translation suppression in cells. Sci Rep 6: 22719.

39. Thandapani P, Song J, Gandin V, Cai Y, Rouleau SG, Garant J-M, Boisvert F-M, Yu Z, Perreault J-P, Topisirovic I, et al. (2015) Aven recognition of RNA G-quadruplexes regulates translation of the mixed lineage leukemia protooncogenes. Elife 4:.

40. Pan L, Li Y, Zhang H-Y, Zheng Y, Liu X-L, Hu Z, Wang Y, Wang J, Cai Y-H, Liu Q, et al. (2017) DHX15 is associated with poor prognosis in acute myeloid leukemia (AML) and regulates cell apoptosis via the NF-kB signaling pathway. Oncotarget 8: 89643–89654.

41. Harashima A, Guettouche T, Barber GN (2010) Phosphorylation of the NFAR proteins by the dsRNA-dependent protein kinase PKR constitutes a novel mechanism of translational regulation and cellular defense. Genes Dev 24: 2640–2653.

42. Wolkowicz UM, Cook AG (2012) NF45 dimerizes with NF90, Zfr and SPNR via a conserved domain that has a nucleotidyltransferase fold. Nucleic Acids Res 40: 9356–9368.

43. Graber T, Baird S, Kao P, Mathews M, Holcik M (2010) NF45 functions as an IRES trans-acting factor that is required for translation of cIAP1 during the unfolded protein response. Cell Death Differ 17: 719–729.

44. Gratacós FM, Brewer G (2010) The role of AUF1 in regulated mRNA decay. Wiley Interdiscip Rev RNA 1: 457–473.

45. Akichika S, Hirano S, Shichino Y, Suzuki T, Nishimasu H, Ishitani R, Sugita A, Hirose Y, Iwasaki S, Nureki O, et al. (2019) Cap-specific terminal N6-methylation of RNA by an RNA polymerase II-associated methyltransferase. Science 363:.

46. Anders S, Huber W (2010) Differential expression analysis for sequence count data. Genome Biology 11: R106.

47. Kassambara A, Mundt F (2017) factoextra: Extract and Visualize the Results of Multivariate Data Analyses.

48. Wickham H, Chang W, Henry L, Pedersen TL, Takahashi K, Wilke C, Woo K, RStudio (2018) ggplot2: Create Elegant Data Visualisations Using the Grammar of Graphics.

