## Supplementary material for "Codon Bias Confers Stability to mRNAs via ILF2 in Humans": Expanded View Figures 1-5

### Expanded View Figure 1

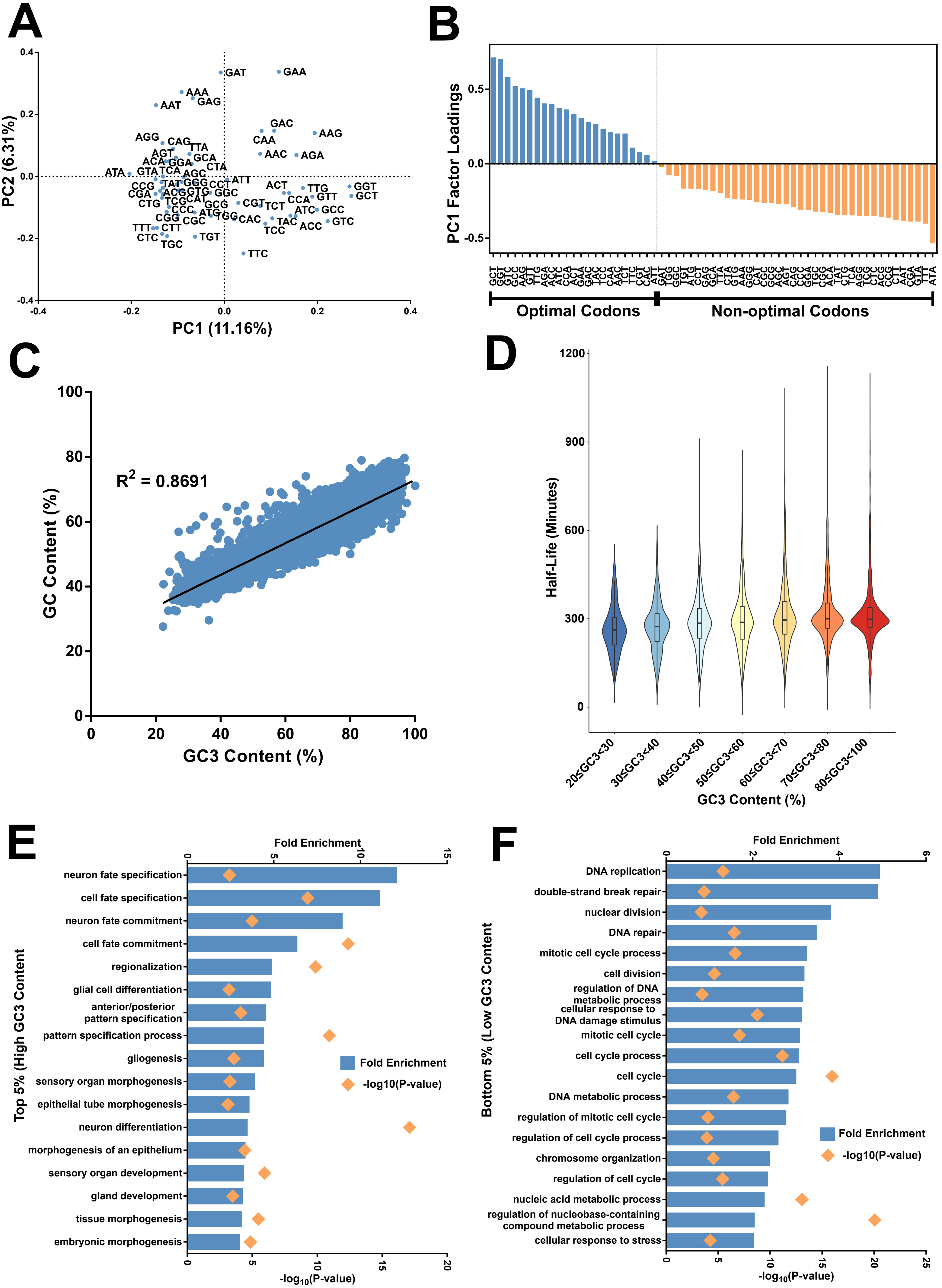

### Expanded View Figure 2

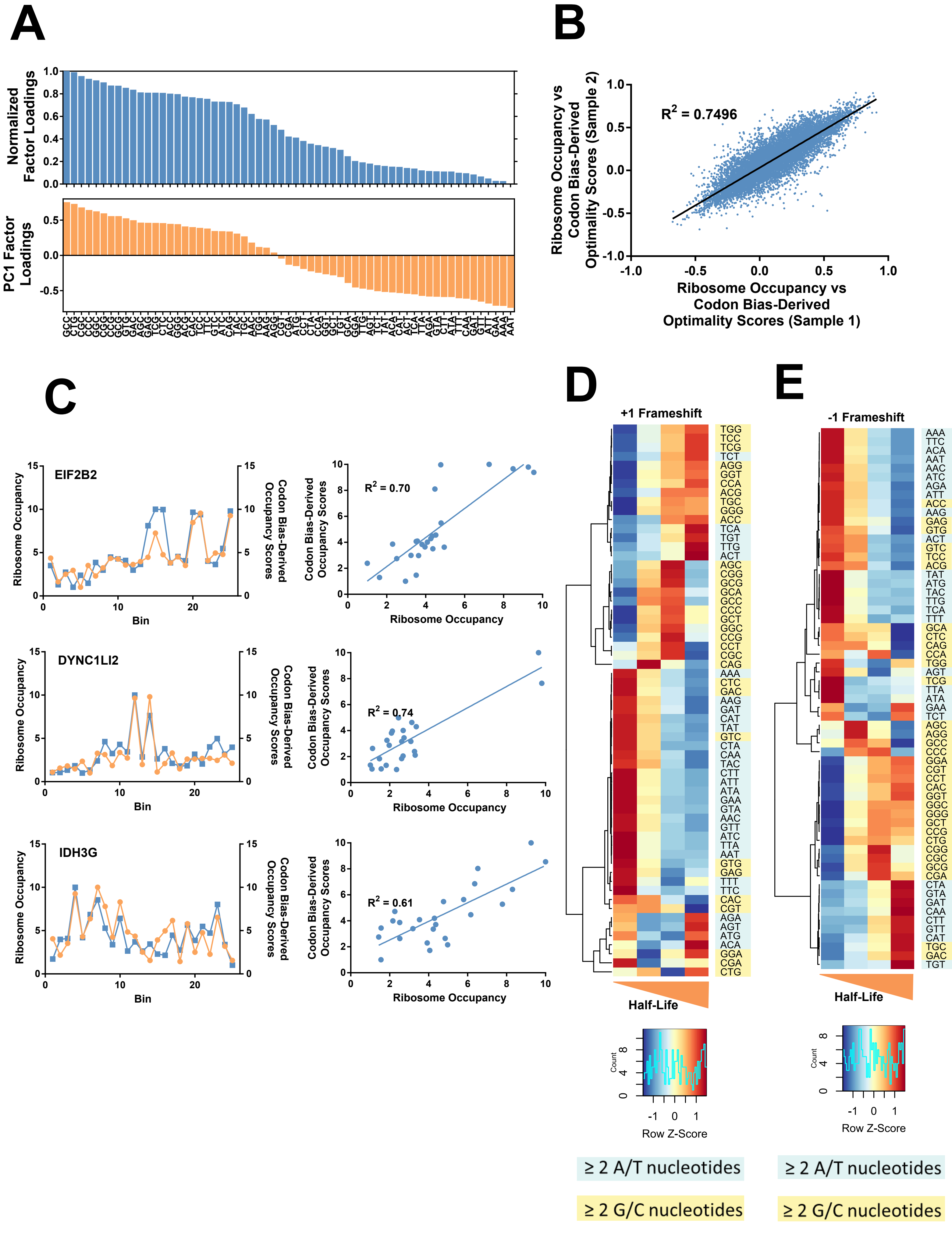

### Expanded View Figure 3

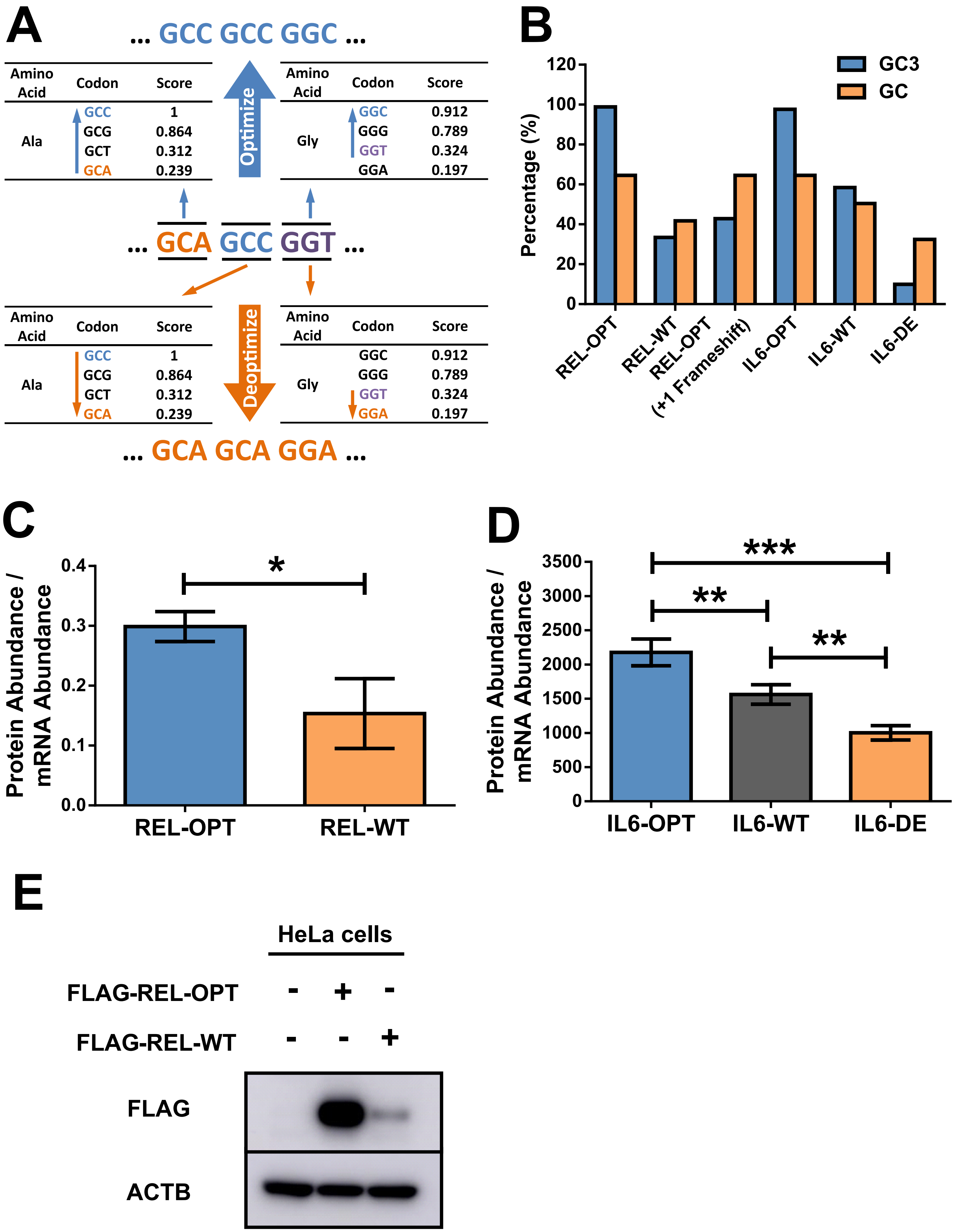

### Expanded View Figure 4

A

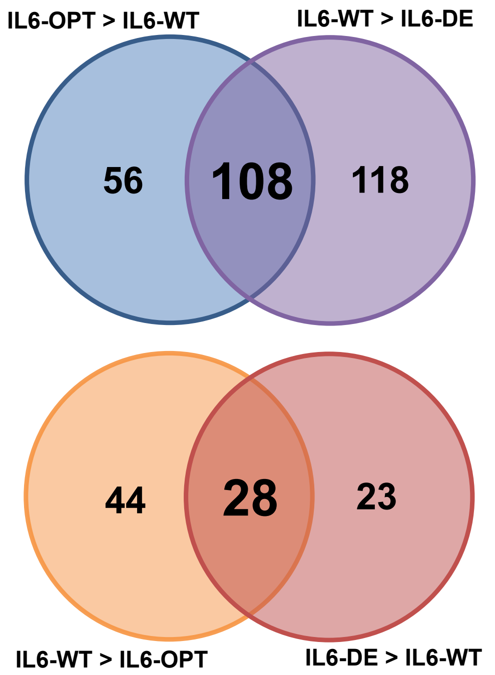


B

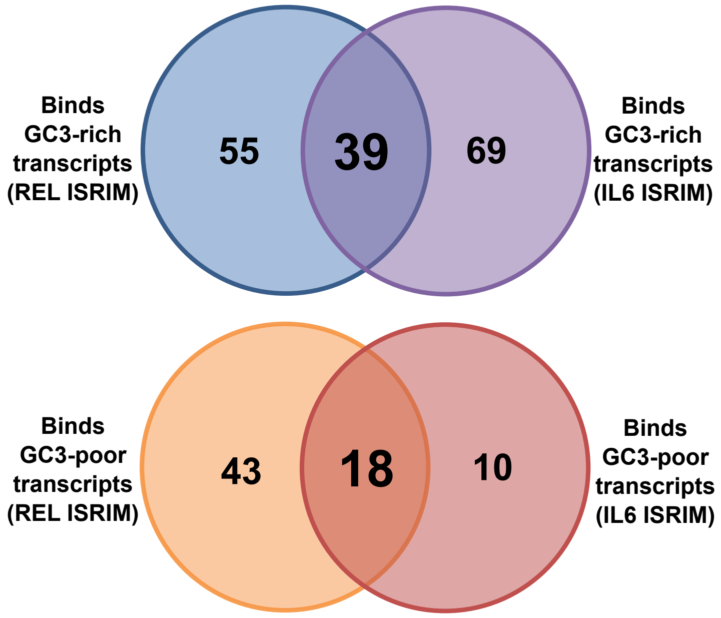

### Expanded View Figure 5

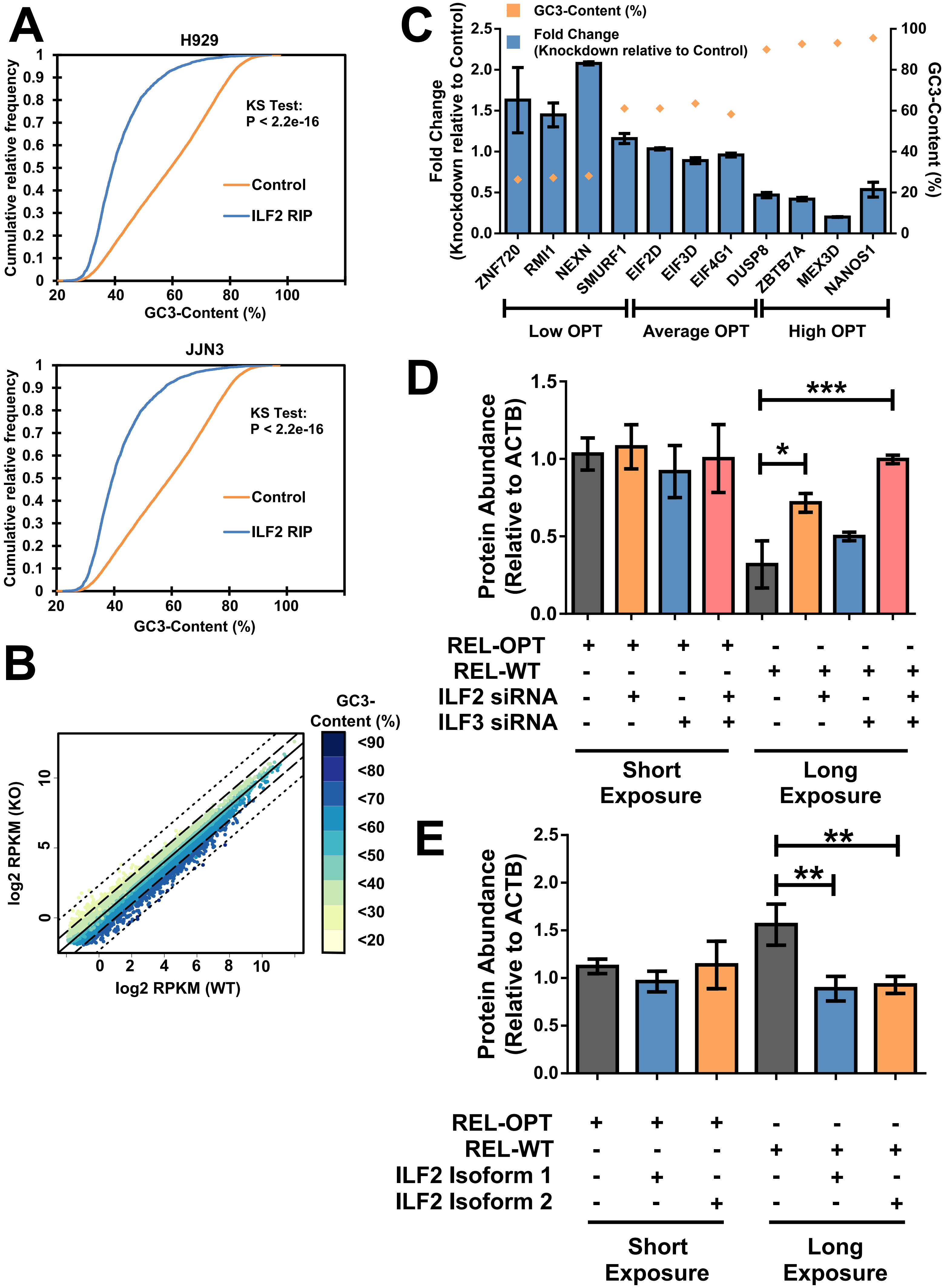
